# Tendon Structure Quantified using Ultrasound Imaging Differs Based on Location and Training Type

**DOI:** 10.1101/393520

**Authors:** Kenton L. Hagan, Todd Hullfish, Ellen Casey, Josh R. Baxter

**Affiliations:** Department of Physical Medicine and Rehabilitation, Perelman School of Medicine, University of Pennsylvania, Philadelphia, Pennsylvania, USA; Department of Orthopaedic Surgery, Perelman School of Medicine, University of Pennsylvania, Philadelphia, Pennsylvania, USA; Department of Physiatry, Hospital for Special Surgery, New York, New York, USA

**Keywords:** Achilles tendon, tendinopathy, ultrasound, running, collagen alignment

## Abstract

Achilles tendinopathy is ten-times more common amongst running athletes compared to age-matched peers. Load induced tendon remodeling and its progression in an at-risk population of developing symptomatic tendinopathy is not well understood. The purpose of this study was to prospectively characterize Achilles and patellar tendon structure in competitive collegiate distance runners over different competitive seasons using quantitative ultrasound imaging. Twenty-two collegiate cross country runners and eleven controls were examined for this study. Longitudinal and cross-sectional ultrasound images of bilateral Achilles and patellar tendons were obtained at the one week prior to start of formal collegiate cross country practices, one week after the conclusion of cross country season, and one week prior to outdoor track and field championships. Collagen organization, mean echogenicity, tendon thickness, and neovascularity were determined using well established image processing techniques. We found that Achilles and patellar tendons respond differently to high-volume running and transitions from one sport season to another, suggesting that tendon structure is sensitive to differences in tendon loading biomechanics. Our findings indicate that Achilles tendon structure in trained runners differ structurally to control tendons but is stable throughout training while patellar tendon structure changes in response to the transition in training volume between cross country and track seasons. These findings expand upon prior reports that some degree of tendon remodeling may act as a protective adaptation for sport specific loading.

**News and Noteworthy:** In this study we prospectively examined the Achilles and patellar tendon structure of distance runners to determine if continued training through multiple seasons elicits tendon remodeling or pathology. We found that Achilles and patellar tendons respond uniquely to the changing loads required during each season. Achilles tendon collagen alignment is mostly stable throughout the competitive cycle, but the patellar tendon undergoes structural changes following the transition from cross-country to track season.

## Introduction

Achilles tendinopathy is a painful degeneration of the tendon that is ten-times more common amongst running athletes compared to age-matched peers (15). During running, the Achilles tendon is cyclically loaded in excess of twelve body weights (14), which is a potential driving factor in tendon remodeling and tendinopathy development in these athletes (5). Distance runners undergo millions of running gait cycles over the course of a competitive season when accounting for training and racing loads. Once symptomatic, Achilles tendinopathy can persist as a chronic problem and limit activity levels in both recreational and competitive athletes.

Tendinopathy manifests at multiple scales, ranging from tissue-level swelling and neovascularization to altered collagen synthesis. Symptomatic tendinopathy can be characterized structurally with increased tendon thickness (19) and decreased collagen alignment (34). Fusiform swelling and increased vascularity are reliably detected using ultrasound imaging (34). Collagen synthesis in the peritendon increases following an intense bout of exercise (19) and prolonged periods of training in novice runners (18), which have not been associated with changes in tendon structure (8). While tendon overload is considered to be a driver of symptomatic tendinopathy (6), eccentric tendon loading often serves as an effective therapy to relieve symptoms (21).

The progression of load induced tendon remodeling in a population at increased risk of developing symptomatic tendinopathy is not well understood (24). Quantitative ultrasound imaging has shown that the Achilles tendons of trained-distance runners are thicker, less organized and, and less echogenic than healthy-young peers despite an absence of clinical symptoms of tendinopathy (11). However, remodeling is not observed in novice runners over the course of 34 weeks despite increased tendon mechanical properties (9). It remains unclear when and how structural changes to tendon occur in response to the repetitive loads of distance running

The purpose of this study was to prospectively quantify Achilles tendon and patellar tendon structure in competitive collegiate distance runners over the course of cross country, indoor track, and outdoor track seasons. Previous studies have shown that trained collegiate runners have structurally different Achilles tendons compared to age-matched peers (11) but the structural response to prolonged training is not well understood. We hypothesized that Achilles tendon structure of distance runners would be quantitatively different compared to healthy-young controls and remain so throughout the rigors of a training season. Furthermore, should clinical signs of tendinopathy develop, we expected to detect tendon structure changes associated with tendinopathy including: increased thickness, decreased collagen alignment, and increased echogenicity (34). We also quantified patellar tendon structure as a ‘control’ tendon, which does not tend to become tendinopathic in distance runners at increased rates compared to the general population. Understanding tendon remodeling in response to prolonged and changing cyclical loads is critical to understanding how tendon disorders progress.

## Methods

### Study Design

Twenty-two collegiate cross country runners (9 females; Age: 19 ± 1.5; Height: 172 ± 7 cm; Weight: 60.4 ± 8 kg) and eleven healthy-young controls (10 females; Age: 24 ± 5.1; Height: 163 ± 7.32 cm; Weight: 59.1 ± 11.6 kg) provided written consent before participating in this IRB approved study. Runners were excluded from this study if they did not continuously remain in full training cycles throughout all three competitive seasons and completed all three scanning sessions. No participants had signs or symptoms of Achilles tendinopathy prior to participating in this study. Ultrasound images and patient reported outcomes were acquired from the runners at the following time points: one week prior to start of formal collegiate cross country practices (S1), one week after the conclusion of the 2017 NCAA cross country season (S2), and one week prior to 2018 NCAA outdoor track and field intercollegiate conference championships (S3). Male and female athletes ran an average of 128 and 80 kilometers each week during the cross country season (S1, S2) and 110 and 70 kilometers each week during the track and field season (S3), respectively. These mileage amounts for the male athletes is consistent with previous of Division I NCAA cross country programs (35).Outdoor track training cycles included one or two, high intensity, anaerobic running sessions per week compared to cross country and indoor track training cycles which had zero or one such sessions. Each of the three visits occurred on a rest day during the participant’s training cycle and at least 12 hours after the most recent running or training session. Furthermore, no participant participated in a designated long run (a run which consumed 20% or more of weekly mileage) or anaerobic based high intensity running workout, or competition in the 24 hours prior to any visit. The VISA-A, a clinical-outcome questionnaire specific to Achilles tendon health (12), was collected in the runner group at each visit in order to determine health and function Achilles tendon. Ultrasound images were acquired in the healthy-young controls during a single measurement session. These controls were recreationally active and had no history of Achilles or patellar tendon injuries or pain; therefore, we did not collect the VISA-A questionnaire on this group.

### Image Acquisition

Ultrasound images of the mid-substance of Achilles and patellar tendons were acquired bilaterally in the longitudinal and transverse planes. Images of the Achilles tendons were acquired while participants lay prone on a treatment table with ankles in the resting position off the end of the table. This position has been used previously to normalize the amount of tension in each subject’s tendon (31). Participants then sat on the treatment table with legs hanging freely (in approximately 90 degrees of knee flexion) while longitudinal and transverse ultrasound images of the mid-substance of the patellar tendon were acquired bilaterally. Images were acquired with an 18 MHz transducer (L18-10L30H-4, SmartUs, TELEMED) with a 3 cm scanning width that was positioned approximately 4 cm proximal to the calcaneal insertion for the Achilles tendon and centrally positioned between the tibial tuberosity and patella for patellar tendon imaging. Scanning parameters were held constant for all subjects over both scanning session (scan parameters: Dynamic Range: 72dB; frequency: 18 MHz; gain: 47 dB). Longitudinal Doppler sequences were also acquired at both the Achilles and patellar tendons in order to quantify tendon neovascularization, which is a clinical indicator of tendon healing and pathology (34). All images were acquired by a single investigator in all sessions and participants as a continuous video and saved as video files.

### Image Analysis

Tendon structure was quantified using objective measures of collagen alignment and tendon thickness. A single frame of each acquisition that clearly showed the mid–substance of the Achilles tendon was selected by the same investigator for all subjects across both imaging sessions. In order to prevent any potential analyses bias, all images were selected and analyzed in a single session in which the trials were randomized and blinded to the investigator. Clinical grading of tendon structure and neovascularization was performed by a fellowship-trained sports medicine physician who was blinded to all other subject data. Tendon structure and neovascularization were both graded (32) on a scale from 0 to 3: ranging from normal (0) to severe symptoms (3).

Collagen alignment was quantified in the longitudinal B-mode ultrasound images using a computational analog for crossed polarizer light imaging (23). This analysis was implemented through a custom-written software that has previously been described and has strong intrarater reliability (10). The banded appearance of tendon results from collagen fascicles appearing hyperechoic and the noncollagenous matrix between the fascicles appearing hypoechoic under US. To quantify the alignment of these bands, a rectangular region of interest of the visible length of the free Achilles tendon was selected, while edges of the tendon were avoided to prevent edge artifacts during analysis. A linear kernel was applied over groups of pixels in the image through a convolution matrix at varying angles between 0 and 180 degrees. The primary direction of each fascicle was calculated as the angle of maximum echo intensity using a power series function. The number of fascicles aligned at each angle was plotted in a histogram. The circular standard deviation (CSD) was calculated by determining the number of collagen fascicles that were not aligned in the primary fiber direction and the magnitude of the difference between their alignment and the mean angle. The CSD and collagen alignment were then, by definition, inversely proportional: the higher the CSD, the less organized the structure of the tendon.

Tendon thickness and echogenicity were calculated using established-quantitative methods (7, 29, 34). Longitudinal thickness of the tendon was measured as the point to point distance between the deep and superficial edges of the tendon. Tendon thickness was calculated by the same blinded investigator using open-source image processing software (29). This measurement has been shown to be both repeatable and reliable in a prior report (34). Average echogenicity was calculated from the same regions of interest that were used to calculate collagen alignment (7).

### Statistical Analysis

Tendon alignment, thickness, and echogenicity were compared between the three study visits within the runner group and between the runner and control groups using a one-way analysis of variance (ANOVA, SigmaStat 4.0, Systat Software, Inc).

Normality (Shapiro-Wilk) and equal variance (Brown-Forsythe) tests were performed on each data set; and when these tests failed, ANOVA based on ranks were performed. In the event that differences between groups were greater than would be expected by chance (P < 0.001), tests that controlled for multiple comparisons were performed. Specifically, Holm-Sidak and Dunn’s methods for parametric and non-parametric tests, respectively. Secondary regression analyses were performed to determine the relationship between collagen alignment (CSD) and mean echogenicity. Six subjects in the runner group dropped out of the study prior to completion due to lower extremity stress fractures or osseous stress reactions which caused the athlete to be removed from the standard running program. Thus, we only included the remaining 16 runners (6 female) for all analyses. Our prior work demonstrated that tendon CSD did not differ between the sexes (11); therefore, we included runners of both sexes in the same analyses.

## Results

Achilles tendinopathy symptoms did not develop in any subjects over the duration of the study period, which was confirmed by clinical ultrasound and VISA-A scores at each testing session (Table 1). During the first session (S1) all runners were found to have qualitatively ‘normal’ structure. Following the first competitive cycle (NCAA cross country season, S2) one runner was found to have ‘mild neovascularization’ under color doppler while all other subjects remained ‘normal.’ The final scanning session that followed the indoor and outdoor track and field seasons (S3) revealed that the Achilles tendon was clinically healthy with no signs of neovascularization or increased echogenicity. No detectable change in VISA-A scores between any sessions was found. Six runners did not complete the scanning session which were all due to non-tendon injuries.

**Table 1.** Subjective measurements of Achilles tendon status

|  | Control | Session 1 | Session 2 | Session 3 |
| --- | --- | --- | --- | --- |
| VISA-A (out of 100) | - | 93 ± 8 | 94 ± 7 | 94 ± 8 |
| Structure (0 – 3) | 0.09 ± 0.30 | 0 ± 0 | 0 ± 0 | 0 ± 0 |
| Neovascularization (0 – 3) | 0 ± 0 | 0.05 ± 0.21 | 0 ± 0 | 0 ± 0 |

Achilles tendon structure differed between runners and controls, and tendon structure changed in runners between the cross-country and track seasons (Figure 1 *top).* Runners demonstrated reduced Achilles tendon alignment (28% increase in CSD) compared to controls, which was similar in scale to increased tendon hypertrophy (13% increase in thickness). While tendon thickness remained stable in the runners throughout cross-country and track season, tendon collagen alignment demonstrated an improvement at the end of the track season compared to the start of the cross-country season (7% decrease in CSD). Mean echogenicity did not differ between the runners and controls or throughout the competitive running seasons.

**Figure 1.**
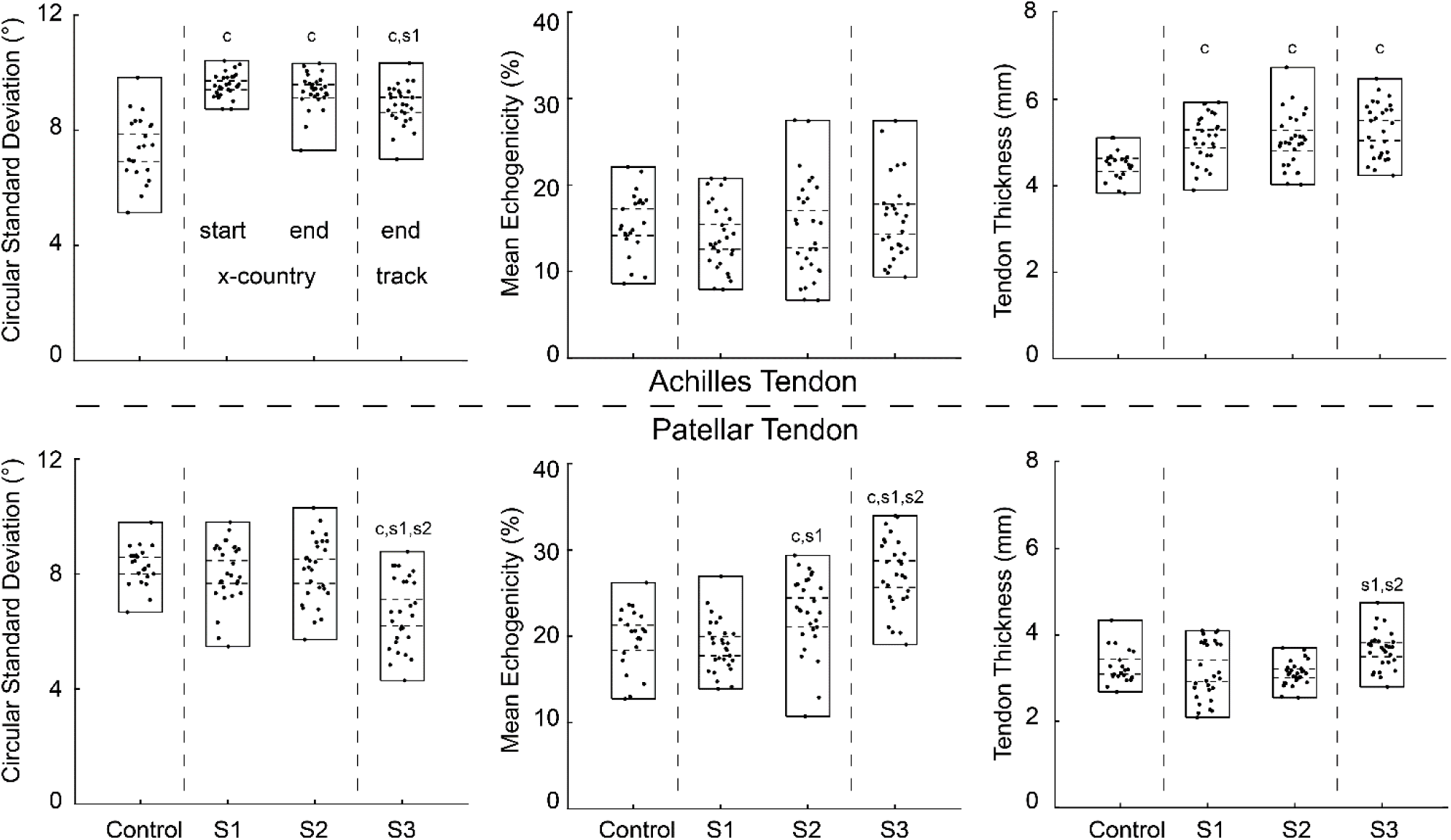
Measures of Achilles tendon (top) and patellar tendon (bottom) circular standard deviation (left), mean echogenicity (middle), and tendon thickness (right) plotted for runners and controls. Runners have three time points corresponding with S1, S2, and S3. The measurement ranges (solid lines) and 95% CI for each group (dashed lines) for each group are plotted. Runners demonstrated reduced Achilles tendon alignment (28% increase in CSD) through S1 and S2 with a 7% improvement at S3. Runners had increased tendon hypertrophy (13% increase in thickness) compared to controls which remained stable. Patellar tendons were not significantly different from controls except at S3 which evealed 17% decrease in CSD, tendon hypertrophy (21% increase in thickness), and 32% increase in average echogenicity.

Patellar tendon structure did not differ between the controls and runners throughout the cross-country season, with the exception of session 2 mean echogenicity (Figure 1 *bottom*). However, at the end of the competitive track season (S3), distance runners underwent collagen alignment (17% decrease in CSD) and tendon hypertrophy (21% increase in thickness). These changes in patellar tendon structure were accompanied by a 32% increase in average echogenicity. Despite observing these structural changes in the patellar tendon, a few solitary blood vessels (Doppler score of 1) was only detected in two runners. All other images of runner patellar tendons demonstrated homogenous echogenicity (echogenicity score of 0) and no neovascularization (Doppler score of 0).

Ultrasound measurements of tendon alignment (CSD) negatively correlated with average tissue echogenicity for both the Achilles and patellar tendons (Figure 2, *P* < 0.001). The Achilles tendon had fundamentally different groupings of these ultrasound measurements between the runners (R^2^ = 0.251) and controls (R^2^ = 0.634), which was explained by the increased CSD values measured in the runners (Figure 1 *top left*). Conversely, the ultrasound measurements of the patellar tendon in the controls and runners demonstrated overlap and were therefore included in a single linear regression model (R^2^ = 0.318).

**Figure 2.**
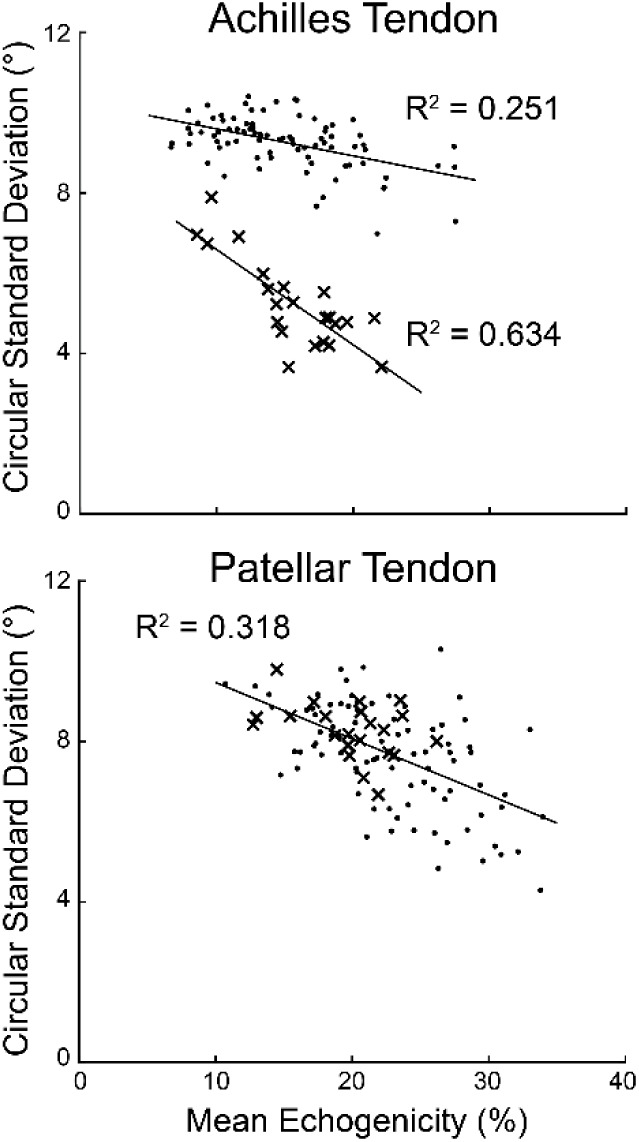
Mean echogenicity was compared with circular standard deviation for runners and controls (top). There was negative correlation between mean echogenicity and circular standard deviation for the runners (P < 0.001). There was overlap between the runners and controls in the patellar tendon and thus a single regression model was used (bottom).

## Discussion

Achilles tendon injuries are ten-times more common amongst running athletes (16), but the effects of continued and high-volume loading on tendon structure remains unclear. Using quantitative ultrasound imaging, we established that trained-distance runners have habituated Achilles tendons in which the collagen alignment deviates less than 7% throughout a competitive season (Figure 1). It is important to note, that these changes were not associated with any symptoms or effects on training and competition. Further, we found that Achilles and patellar tendons respond differently to high-volume running and transitions from one sport season to another, suggesting that tendon structure is sensitive to differences in tendon loading biomechanics. Although ultrasound measurements of collagen alignment (CSD) correlated with tissue echogenicity (Figure 2), quantifying collagen alignment appears to have greater sensitivity to changes in tendon structure and may be a preferable image processing technique for studying subtle changes to tendon structure in response to prolonged loads.

Sport specific tendon adaptations have been well documented in athletes of various competitive levels (6, 9, 21, 33). A previous study in distance runners showed small amounts of Achilles tendon hypertrophy during a competitive running season (33). Similarly, analysis of rotator cuff tendons in elite swimmers show darker and thicker tendons indicative of tendon habituation to long-term training and competition (26). Chronic cyclical loading induces structural tendon changes which appear to be a protective adaptation developed to protect trained runners against injury risk (11, 21, 27). Thicker tendons undergo less strain when exposed to cyclic loads compared to normal tendon (27). Additionally, tendon remodeling also has a role in modulating the performance of a tendon. Distance runners have decreased collagen alignment and echogenicity compared to healthy controls (11). Furthermore, specific loading types play a significant role in tendon remodeling as sprinters have been shown to have stiffer tendons when compared to distance runners and healthy controls (2, 9). Our findings indicate that Achilles tendon structure does undergo some realignment (as measured by CSD) after the start of the competitive running season but remains stable throughout multiple competitive cycles while the patellar tendon becomes increasingly more organized at the conclusion of a collegiate running season.

Exercise has been shown to increase collagen synthesis in humans (18, 19); however, the clinical effects of this increased synthesis have not been made, nor have they been directly linked to tendon remodeling. Furthermore, these studies have been limited to short term alterations in activity levels within previously healthy individuals. It is unclear if highly-trained distance runners maintain this increased collagen synthesis given their habituation to prolonged tendon loading. The magnitude and frequency of tendon loading also impacts tendon remodeling (1, 3). The varying mechanical adaptations elicited as a result of varying loading frequencies appear to agree with differences previously observed between long distance runners and sprinters (2). To maximize the velocity-to-energy ratio, trained runners increase tendon loading to in order to efficiently store and return elastic energy. For example, the metabolic cost of distance running has been correlation with smaller Achilles tendon moment arms, which highlights the importance of increased tendon loading during running (22, 30).

Tendon habituation in response to prolonged loading may serve as a protective mechanism against overuse injuries. Achilles tendon habituation or remodeling does not occur in novice runners in a low intensity program over 34 weeks (9). The present study demonstrates that over the course of a competitive season, particularly later in the competitive cycles of indoor and outdoor track, previously habituated Achilles and patellar tendons undergo detectable amounts of collagen realignment (Figure 1). It appears that training alterations for track competitions when compared to cross country or pre-season training induces tendon remodeling. Rapid changes in activity level may increase the development of tendinopathy (4), suggesting that the loading thresholds for tendon remodeling and pathologic development depend on the history of training. This study cohort maintained a regimented training routine and did not develop tendinopathy despite being at a nearly ten-fold higher injury risk (16). It remains unclear when these structural, and seemingly permanent, changes to the Achilles tendon occurred. If indeed tendon remodeling to load occurs during adolescence (32), then these adaptations may serve as a protective mechanism against increased tendon loading later in life.

Several limitations should be considered when interpreting these results. First, quantitative ultrasound images were acquired and measured by a single investigator; but these measurements have been shown to be repeatable and reliable between and within investigators (10), and the investigator was blinded to the image identifier during analysis. Additionally, a single-fellowship trained sports medicine physician interpreted all images for neovascularity. All participants in this study were part of the same collegiate cross-country team and underwent similar training regimens, but each athlete’s weekly mileage varied and factors such as competition schedules, diet, running mechanics, medication exposure, and genetics were unable to be controlled. At the outset of this study, we anticipated that some runners would develop symptomatic Achilles tendinopathy. No symptoms were detected throughout this study, which may have been a result of individualized training programs based on running history, effective coaching, and athletic training. Tendon mechanical properties were not measured in this study; however, prior reports have linked tendon mechanics with alignment (16, 17) and echogenicity (7).

In conclusion, Achilles tendons in highly-trained distance runners are structurally different than recreati onally active peers and appear to be structurally stable during a single competitive season, which suggests tendon habituation to sport-specific loading. However, these same runners underwent quantitative structural changes of both the Achilles and patellar tendons following a transition in training volume without the development of symptomatic tendinopathy. These findings expand upon prior reports that some degree of tendon remodeling may act as a protective adaptation for sport specific loading. It remains unclear if changes in the structure of the Achilles or patellar tendons are precursors of future tendon dysfunction; however, the training rigors of highly-trained distance runners at the collegiate level does not appear to cause deleterious effects on the tendon, and in fact may produce further protective changes. Further understanding the process of tendon remodeling in both healthy and pathologic mechanisms is essential to improving diagnosis and treatment of tendon disorders.

## Acknowledgements

We would like to thank Annelise Slater and Neza Stefanic for assistance in data collection and Jamel Jones and Dr. Alexis Tingan for assistance in subject recruitment. This work was supported by the Thomas B. McCabe and Jeannette E. Laws McCabe Fund.

This study was performed at The University of Pennsylvania.

